# Notch-Mediated Polarity Decisions in Mechanosensory Hair Cells

**DOI:** 10.1101/480798

**Authors:** A. Jacobo, A. Dasgupta, A. Erzberger, K. Siletti, A. J. Hudspeth

## Abstract

The development of mechanosensory epithelia, such as those of the auditory and vestibular systems, results in the precise orientation of mechanosensory hair cells and consequently directional sensitivity. After division of a precursor cell in the zebrafish’s lateral line, the daughter hair cells differentiate with opposite mechanical sensitivity. Through a combination of theoretical and experimental approaches, we show that Notch1a-mediated lateral inhibition produces a bistable switch that reliably gives rise to cell pairs of opposite polarity. Using our mathematical model of the process, we predict the outcome of several genetic and chemical alterations to the system, which we then confirm experimentally. We show that Notch1a downregulates the expression of Emx2, a transcription factor known to be involved in polarity specification, and acts in parallel with the planar-cell-polarity system to determine the orientation of hair bundles. By analyzing the effect of simultaneous genetic perturbations to Notch1a and Emx2 we infer that the gene-regulatory network determining cell polarity includes undiscovered polarity effectors.

## Introduction

Notch-Delta signaling regulates diverse processes in development, regeneration, and disease. In many contexts signaling through this pathway creates cellular patterns through suppressive cell-cell interactions, in which the concentrations of signals in interacting cells diverge over time, leading to distinct cellular fates (*1*–*3*). We have discovered a novel role for Notch-mediated lateral inhibition in determining the polarity of sensory hair cells during development and regeneration.

By transducing mechanical stimuli into electrical signals, hair cells underlie our senses of hearing and balance (*4*). The polarity of a hair cell is morphologically determined by its hair bundle, which protrudes from the cell’s apical surface and confers directional sensitivity to mechanical stimuli (*5*). Although signaling by the core planar-cell-polarity (PCP) system aligns hair bundles with the polarity axis of a sensory organ(*6*, *7*), some hair cells reverse their polarity with respect to the PCP axis, rendering them optimally responsive to mechanical stimuli from the opposite direction (*8*, *9*). The transcription factor Emx2 is specifically expressed in hair cells with reversed polarity and participates in the specification of that fate (*10*), but the mechanism that establishes the Emx2 identity of a hair cell and effects polarity reversal remains unknown.

Using the lateral line of the zebrafish as a model system, we have studied how the correct number of oppositely polarized hair cells is established during the formation of mechanosensory epithelia. The lateral line consists of a series of sensory organs called neuromasts that sense water movements (Figure 1A) (*9*). Hair cells in the neuromast develop in pairs from a progenitor cell, with the two daughter cells bearing oppositely polarized hair bundles (*9*). One cell is sensitive to caudad water motion—flow toward the tail—whereas the sibling cell responds to rostrad movement—flow toward the head (Figure 1B,C). The expression of the core PCP protein Vangl2 is indistinguishable in the two cells (Figure 1D,E), but only caudad-polarized cells express Emx2 (Figure 1F,G) (*10*). Here we demonstrate that lateral inhibition mediated by Notch-Delta signaling dictates the binary fate decision involving Emx2 that underlies polarity patterning.

**Figure 1:**
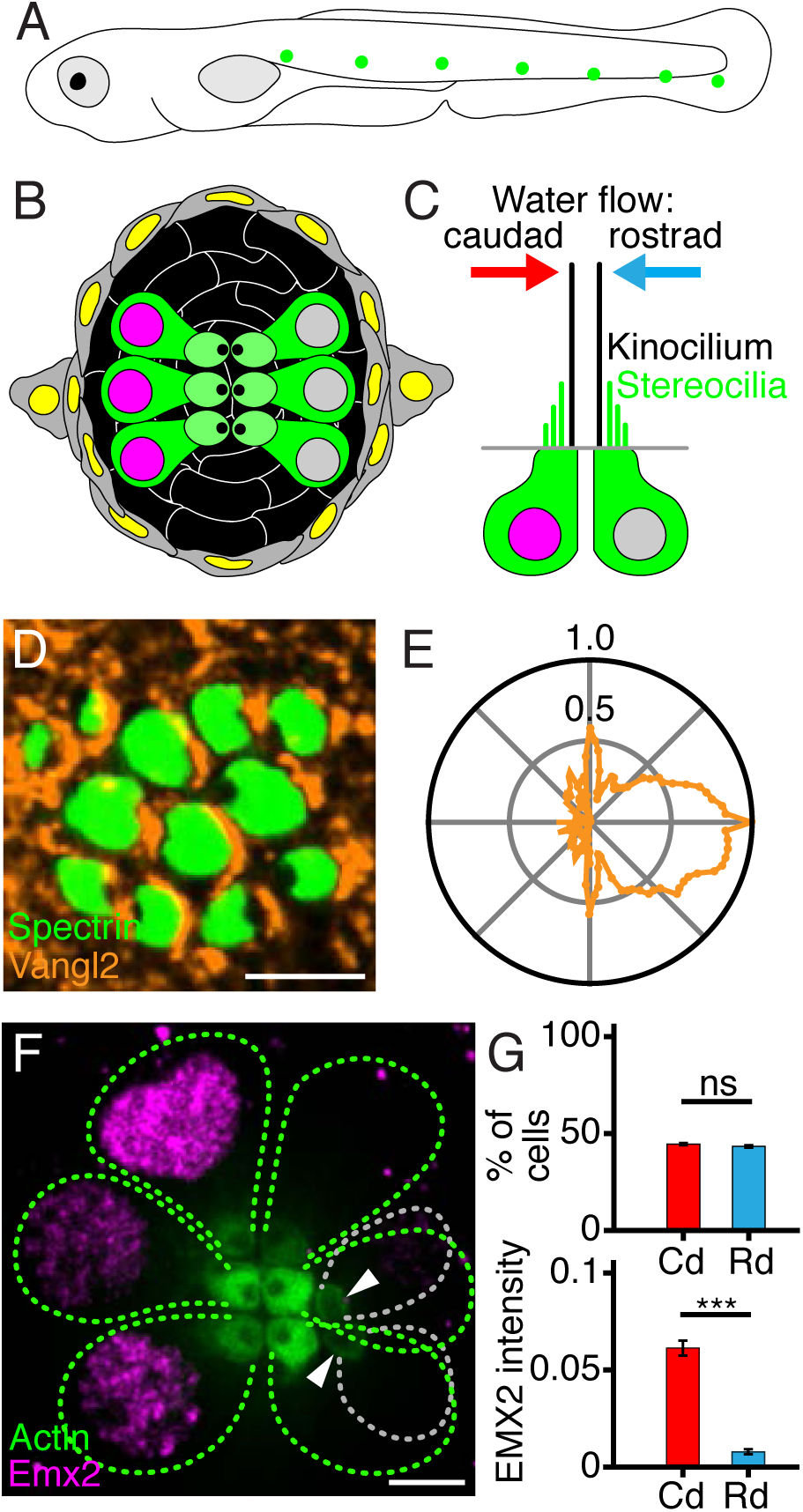
Organization of neuromasts in the zebrafish’s lateral line. (**A**) A schematic drawing of a 4 dpf larva depicts neuromasts (green) of the posterior lateral line. Additional neuromasts (not shown) adorn the anterior portion of the larva. (**B**) A diagramatic surface view of a single neuromast shows hair cells (green) separated by supporting cells (black) and surrounded by mantle cells (gray). (**C**) A schematic section depicts the sensitivity of oppositely oriented hair cells to water flow. (**D**) In a micrograph of the upper surface of a neuromast, spectrin (green) marks the cuticular plates of hair cells. Immunolabeling reveals that the core PCP protein Vangl2 (orange) occurs consistently at the posterior cellular boundaries. (**E**) A polar plot of average Vangl2 labeling from 502 hair cells quantitates the effect in panel **D**. This and each of the subsequent polar plots represents the average intensity of labeling, normalized to the maximum value, as a function of angular position with respect to the cells’ centers (Supplementary Material). (**F**) In a developing neuromast, Emx2 (magenta) is expressed in the nuclei of three mature hair cells (green dashed lines on the left). In the distribution of actin-GFP (green) at the cellular apices, the dark spots locate the kinocilia and indicate that those cells are sensitive to caudad stimulation. Three other mature hair cells (green dashed lines on the right) have an opposite orientation. The apices of two immature hair cells (gray dashed lines) are indicated by arrowheads. (**G**) The polarities of 81 hair cells from 18 neuromasts (top) are equally divided between caudad (Cd) and rostrad (Rd). The former cells express significantly higher levels of Emx2 (bottom). Scale bars, 3 µm. Means ± SDs; *** = p < 0.001; ns = not significant.

## Results

### Notch-mediated lateral inhibition and polarity reversal

Pharmacological or genetic interference with Notch signaling yields neuromasts with a polarization bias of their hair cells (*11*, *12*). This observation raises the possibility of an interaction between the Notch pathway and Emx2 in the establishment of polarity reversals. To explore this hypothesis, we developed a mathematical model of Notch signaling between a pair of nascent hair cells based on the framework in Boareto *et al*. (Figure 2A; Supplementary Information) (*13*). We simulated the concentration dynamics of ligands and receptors on the cellular surfaces as well as the concentration of Notch intracellular domain (NICD), which is released by proteolytic cleavage after the interaction of Notch with the lig- and Delta and acts as a transcriptional co-activator. We used realistic kinetic rates for binding, unbinding, receptor cleavage, and nuclear translocation (Supplementary Table S2; Supplementary Material) (*13*, *14*).

**Figure 2:**
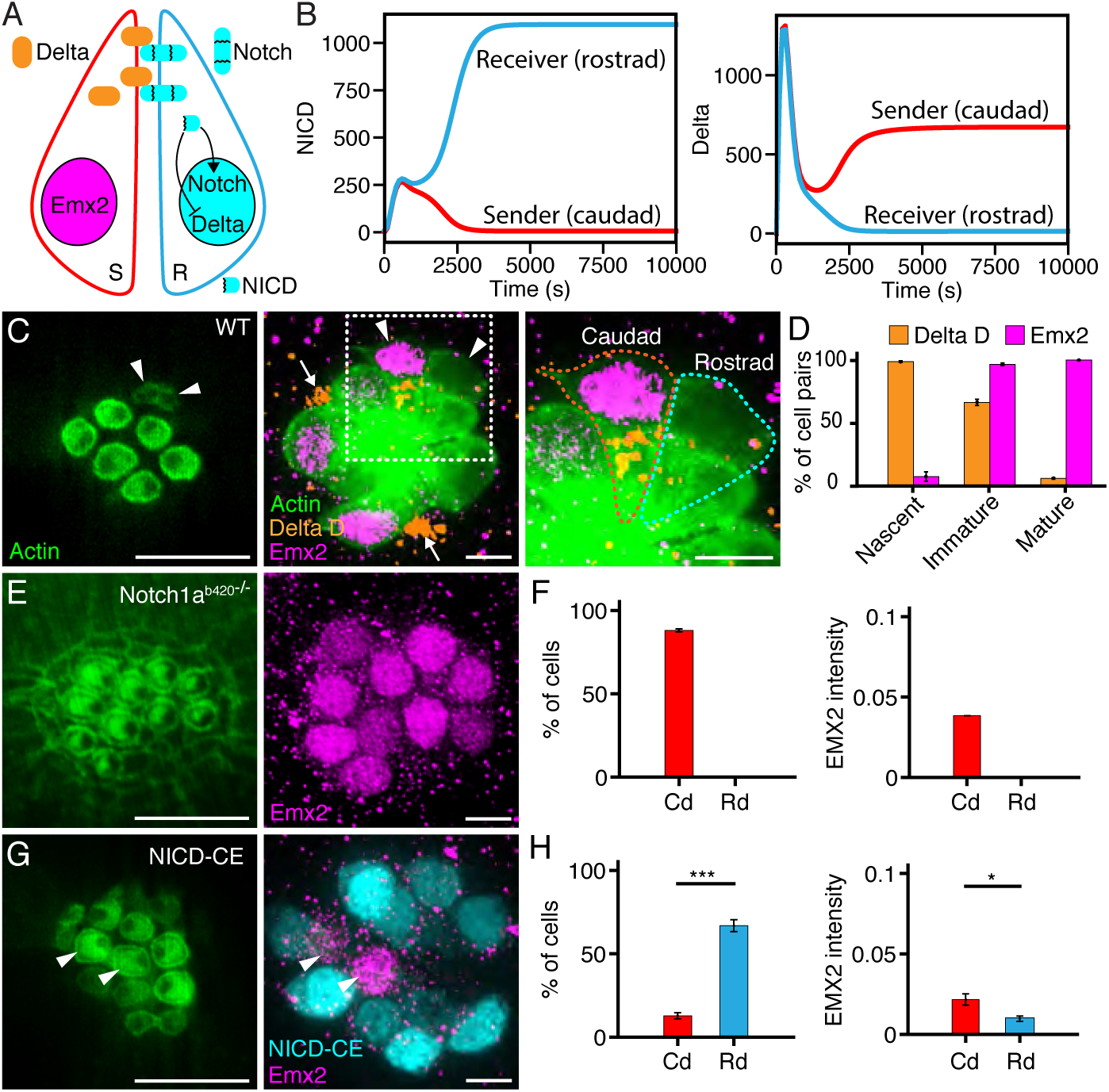
Notch-mediated lateral inhibition and hair-cell polarity. (**A**) A schematic representation of our Notch signaling model shows a pair of nascent hair cells where the Delta ligand (orange) expressed on one cell causes cleavage of Notch receptor (blue) on the other. Upon entering the nucleus, NICD upregulates Notch expression, downregulates Delta expression, and conduces to a receiver phenotype (R). The sister cell evolves in the opposite direction to a sender fate (S). (**B**) Representative simulations portray the concentrations of NICD (left) and Delta (right) in the receiver cell—which becomes rostrad-polarized—and the sender cell—which becomes caudad-polarized. (**C**) An apical view of a 4 dpf wild-type neuromast (left) reveals three mature pairs of hair cells and one immature pair (arrowheads). A view at the nuclear level (center) shows immunolabeling of the same neuromast for Emx2 (magenta) and Delta D (orange). An enlargement of the boxed area (right) includes outlines of the two immature hair cells. Hair-cell progenitors also express Delta D (arrows). (**D**) Delta D is expressed in at least one hair cell of each pair at earlier stages of development than Emx2 (51 nascent pairs, 101 immature pairs, 259 mature pairs). (**E**) In a Notch1a^b420-/−^mutant, hair cells show a caudad polarization (left) and uniformly express Emx2 (right). (**F**) These effects are consistent in 536 hair cells from 57 neuromasts. (**G**) In an NICD-CE transgenic animal, most hair cells are sensitive to rostrad stimulation (left). Occasional hair cells (arrowheads) fail to express NICD-CE; they instead contain Emx2 (magenta, right) and adopt a caudad sensitivity. (**H**) The effect is quantified for 68 cells from 21 neuromasts. Scale bars, 5 µm. Means ± SDs; *** = p < 0.001; * = p < 0.02.

*Trans*-activation of the Notch receptor inhibits the expression of its ligand Delta, creating a negative feedback loop (Figure 2A). Although our simulations started with similar concentrations of NICD in each of the two cells, feedback therefore amplified the initial stochastic differences of NICD expression in the nascent hair-cell pair and broke the initial symmetry (Figure 2A,B). One of the cells evolved to a low-Notch and high-Delta phenotype called the sender state (*14*). That cell suppressed the expression of Delta in its sister, which then achieved a high-Notch phenotype termed the receiver state (Figure 2B). Given that the expression of Emx2 also shows distinct levels in sister cells (Figure 1F,G), we propose that Notch signaling regulates the expression of Emx2, giving rise to a pair of hair cells of which one expresses a high and the other a low concentration of this protein. This arrangement reliably creates one cell with caudad orientation and another with rostrad orientation (Supplementary Material).

To test our model, we investigated the expression of the Notch ligands Delta C and Delta D in regenerating neuromasts of zebrafish larvae at four days post-fertilization (4 dpf). Nascent hair cells that had recently originated from a progenitor cell expressed higher levels of Delta D than mature hair cells with fully formed hair bundles. These nascent cells neither possessed an apical surface nor expressed Emx2. In slightly older but still immature pairs of hair cells, defined as those with underdeveloped apical surfaces, Delta D occurred at the contact area between the cells and was sometimes asymmetrically enriched in the cell that expressed Emx2 (Figure 2C; Supplementary Figure 1A-C; Supplementary Movies 1-3). Furthermore, we quantified the expression of Delta D and Emx2 in nascent, immature, and mature haircell pairs and found that Delta D expression preceded that of Emx2 (Figure 2D). This finding suggests that Notch signaling anticipates the expression of Emx2 during polarity specification. During the maturation of a hair-cell pair the sender cell, with the higher Delta concentration and significant Emx2, has a low Notch concentration. The results suggest that Notch downregulates Emx2 expression and support the hypothesis that Notch-mediated lateral inhibition breaks the symmetry between the two nascent hair cells and defines their Emx2 status. Moreover, the observations imply that the sender cell is destined to develop caudad polarization and the receiver cell to assume rostrad polarization.

### Perturbations of Notch gene expression

If Emx2 expression is regulated through Notch signaling, our model provides two predictions for experimental testing. First, in the neuromasts of a Notch-knockout animal, all hair cells should be Emx2-positive and display caudad sensitivity. And second, constitutive expression of NICD in hair cells should yield two high-Notch, low-Emx2 cells with a rostrad polarity.

To experimentally test the first prediction of our model, we analyzed neuromasts from larvae deficient in functional Notch proteins. In homozygous *Notch1a^b420-/−^* mutants (*15*) we found that all hair cells were Emx2-positive and displayed a caudad polarity (Figure 2E,F; Supplementary Figure 2A). By contrast, homozygous *Notch2^el515-/−^* mutants (*16*) showed no polarity bias: either Notch2 is not involved in polarity regulation or Notch1a compensates for its absence (Supplementary Figure 2B). We additionally examined neuromasts from *deltaA*^−/−^, *deltaC*^−/−^, and *deltaD*^−/-^ mutants (*17*, *18*) and observed no bias in hair-cell polarity (Supplementary Figure 2B). Although the asymmetrical localization of Delta D in immature hair-cell pairs suggests that the protein participates in lateral inhibition, the lack of polarity bias in Delta D mutants points to compensation through functional redundancy among the three ligands.

To test our second prediction we developed a transgenic zebrafish line that constitutively expresses myc-tagged NICD (NICD-CE) specifically in hair cells (Supplementary Table S1; Supplementary Material). As reported previously (*12*), hair cells of these transgenic larvae display a strong rostrad bias compared to those in control animals (Figures 1G and 2G). To investigate the mechanism behind this bias, we used specific antibodies to analyze the Emx2 level in NICD-CE hair cells. Because expression was sometimes mosaic, we restricted our analysis to hair cells expressing NICD-CE as indicated by anti-myc immunolabeling (Figure 2G). Quantification of nuclear expression of Emx2 in both control and NICD-CE hair cells revealed that NICD decreased the expression of Emx2, resulting in rostrad polarity (Figure 2H). We corroborated this result with another transgenic line that induced endogenous production of NICD through cytokine signaling (Supplementary Figure 3 A,B; Table S1; Supplementary Material). Taken together, measurements of polarity distribution and Emx2 expression levels in mutant and transgenic lines show that Notch regulates polarity reversal.

### Notch-Delta signaling and global PCP signals

Because Notch signaling determines Emx2 expression in developing pairs of hair cells, we wondered how the Notch pathway interacts with cell-intrinsic polarity factors and with global PCP signals to determine the orientation of hair bundles. In *trilobite* mutants that lack functional Vangl2 protein, hair bundles are randomly oriented (*7*). Because Emx2 is nevertheless expressed in half of the *Vangl2* ^−/−^ mutant hair cells (Supplementary Figure 4) (*10*), though, a Notch-based lateral-inhibition circuit remains intact in *Vangl2* ^−/−^ mutants and might act in parallel to the PCP system.

The shape and orientation of the hair bundle is determined by the localization of cell-intrinsic polarity factors including the heterotrimeric G-protein Gnai (*19, 20*). Reading cues from the global PCP pathway, Gnai and the associated proteins form a crescent-shaped domain at the apical surface of a developing mammalian hair cell. These factors are implicated in relocalizing the kinocilium from its initial position at the center of the apical surface to the edge where the Gnai crescent lies (*19*, *20*). This process creates a cell-intrinsic polarity axis that defines the orientation of the hair bundle. Whereas at least three isoforms of Gnai occur in the zebrafish genome, RNA-seq data indicate that Gnai1 is highly enriched in lateral-line hair cell (*21*). Regardless of the cellular polarity, Gnai1 adorned the apical surface of each wild-type hair cell as a crescent adjacent to the kinocilium (Figure 3A). Even in immature hair cells with an underdeveloped apical surface, Gnai was asymmetrically distributed, suggesting that Gnai plays a role similar to that of its mammalian homolog.

**Figure 3:**
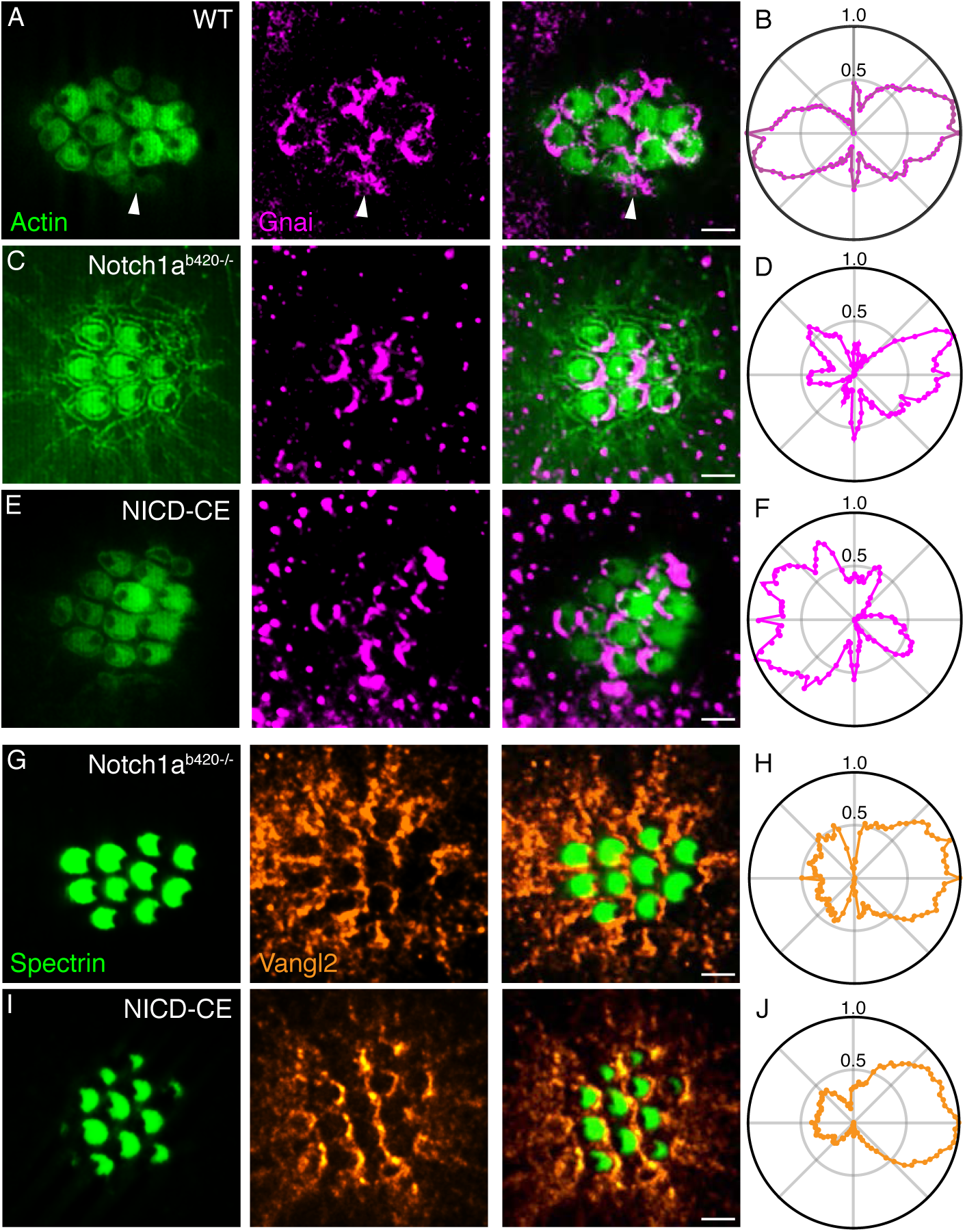
Parallel action of Notch and PCP. (**A**) In an apical view of a wild-type neuromast, hair-cell polarity is delineated by actin-GFP (green, left) and Gnai is immunolabeled (magenta, center). As shown in the merged view (right), Gnai already displays a polarized location in immature hair cells (arrowheads). (**B**) In the average radial intensity profile of Gnai from 239 hair cells of both polarities, caudad and rostrad cells create symmetric lobes on the two halves of the apical surface. (**C**) Notch1a^b420-/−^ hair cells display a consistently caudad orientation and crescents of Gnai at their posterior boundaries. (**D**) A polar plot from 81 hair cells confirms the asymmetrical distribution of Gnai. (**E**) NICD-CE hair cells are consistently sensitive to rostrad stimulation and bear Gnai at their anterior edges.A polar plot from 35 hair cells confirms the asymmetry of Gnai distribution. (**G**) Immunolabeling for spectrin (green, left) shows that Notch1a^b420-/−^ hair cells have caudad sensitivity; labeling for Vangl2 (orange, center) displays its distribution at the posterior edges of the cells. (**H**) A polar plot from 155 hair cells confirms the asymmetrical distribution of Vangl2. (**I**) Although NICD-CE hair cells are consistently sensitive to rostrad stimulation, they also bear Vangl2 at their posterior edges. (**J**) A polar plot from 235 hair cells confirms the asymmetry. Scale bars, 2 µm.

We quantified the average intensity of Gnai1 expression across the apical surfaces of hair cells and observed a mirror-symmetric distribution (Figure 3B). This opposing pattern of localization was lost in *Notch1a^b420-/−^* mutants (Figure 3C,D) and NICD-CE larvae (Figure 3E,F), in which the distribution of Gnai1 nonetheless coincided with the orientation of the hair bundles. The mirror symmetry of the Gnai1 distribution was also absent from *Emx2^−/−^* mutants and larvae constitutively expressing Emx2 (Emx2-CE) (Supplementary Figure 4 A,B; Table S1) (*10*). Notch and Emx2 therefore act upstream of the cell-intrinsic determinants of hair-cell polarity and guide their localization during development.

To further investigate whether core PCP components were affected in Notch-deficient and NICD-CE larvae, we analyzed the localization of Vangl2 protein. In wild-type larvae, Vangl2 was enriched at the posterior edge of a hair cell’s apical surface irrespective of the bundle’s orientation (Figure 1D,E). This expression pattern was maintained in the neuromasts of both *Notch1a^b420-/−^* mutants (Figure 3G,H) and NICD-CE larvae (Figure 3I,J). A similar pattern also occurred in *Emx2^−/−^* mutants and Emx2-CE transgenic larvae (Supplementary Figure 4C,D). Because the distribution of Vangl2 was unaffected by a deficit or excess of Notch function, Notch signaling operates in parallel to the core PCP pathway.

### Interaction of Notch and Emx2 with downstream targets

Emx2 is necessary and sufficient to determine the polarity of hair cells (*10*). Because our observations suggested that Emx2 functions downstream of Notch, expression of both NICD and Emx2 would be expected to yield hair cells with caudad sensitivity similar to those in Emx2-expressing larvae. We therefore developed a line of transgenic zebrafish that constitutively expressed both Emx2 and NICD within hair cells (Figure 4A; Supplementary Figure 5; Supplementary Movie 6; Supplementary Table S1). Surprisingly, hair cells displayed a rostrad polarity bias similar to that characteristic of NICD-CE alone (Figure 4B). This result implies that Emx2 and Notch interact in a more complex manner than the simple downregulation of Emx2 by Notch. We found that in Emx2-CE hair cells the endogenous Emx2 retained a pattern of high expression in one sister cell and low expression in the other (Supplementary Figure 6), suggesting that there is no feedback regulation of the Notch pathway through Emx2.

**Figure 4:**
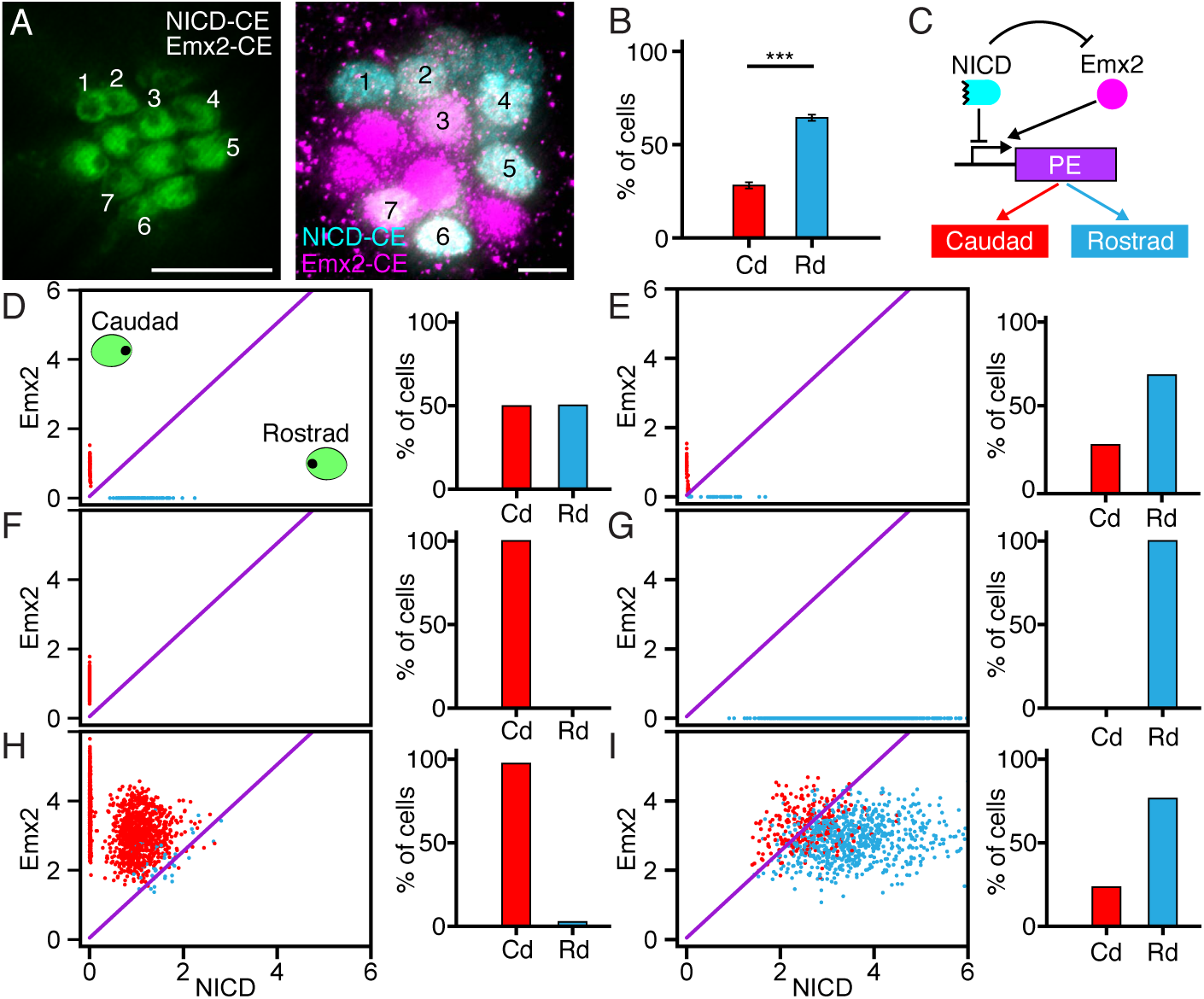
Figure 4. Modeling of the regulatory network underlying hair-cell polarity. (**A**) In an apical view, numerals mark the hair cells in a larva constitutively expressing both NICD (cyan) and Emx2 (magenta). (**B**) In larvae expressing both proteins, 334 hair cells from 50 neuromasts display a strong bias toward rostrad polarity. (**C**) In the proposed regulatory network, Notch and Emx2 control hair-cell polarity through a downstream polarity effector (PE), which might be a single protein or a more complex system. (**D**-**I**) The panels display the steady state expression of Emx2 and NICD for multiple simulations of hair-cell pairs of (**D**) wild-type, (**E**) DAPT-treated, (**F**) Notch1a^−/−^, NICD-CE, (**H**) Emx2-CE, and (**I**) NICD-CE/Emx2-CE neuromasts. In each panel the oblique purple line demarcates the threshold for switching of the polarity effector. Scale bars, 5 µm. Means ± SDs; *** = p < 0.001.

To explain these results, we extended our mathematical model to include an additional hypothesis: NICD and Emx2 compete for the regulation of genes that encode one or more polarity effectors. We propose that Emx2 activates a polarity effector and NICD inhibits it (Figure 4C; Supplementary Information). By symmetry, the model would also operate if these roles were reversed. The concentration of the polarity effector establishes cellular polarity: high concentrations yield caudad-polarized cells whereas low concentrations produce cells of rostrad sensitivity. Under these assumptions our model reproduced the results of all the experimental perturbations that we had performed, including the constitutive expression of both NICD and Emx2 (Figure 4D-I). By including noise in the model we were also able to quantitatively reproduce the polarity biases observed experimentally (Figure 2E-H; Figure 4A,B; Supplementary Figure 8A,B).

In wild-type neuromasts the lateral-inhibition mechanism splits sister cells into two states, low-Notch/high-Emx2 and high-Notch/low-Emx2. Cells with low Notch and high Emx2 concentrations drive the expression of the polarity effector above a certain threshold and become caudad-polarized, whereas in cells with high Notch and low Emx2 concentrations the expression of the polarity factor is repressed and they become rostrad-polarized (Figures 1G and 4D; Supplementary Figure 8).

The first indication that Notch plays a role in regulating polarity reversals was the observation that treatment with *N*-[*N*-(3,5-difluorophenacetyl-L-alanyl)]-S-phenylglycine *t*-butyl ester (DAPT), a *γ*-secretase inhibitor that prevents the liberation of NICD, produced neuromasts with a rostrad bias (Supplementary Figure 9) (*11*, *22*). Because DAPT blocks Notch activity, treated neuromasts might be expected to show a caudad bias similar to that of *Notch1a^b420-/−^* mutants. The seemingly paradoxical result can be explained in light of our model by assuming that DAPT inhibits Notch cleavage incompletely. In this case the bistable circuit that generates high- and low-NICD states fails for many nascent hair-cell pairs. Both cells adopt an intermediate level of NICD that nonetheless suffices to repress Emx2 and then to drive the expression of the polarity effector to sub-threshold levels (Figure 4E; Supplementary Figure 9; Supplementary Information). If the activity of Notch were inhibited more completely, our model predicts that cells would show a caudad bias. Although the regime of low NICD expression was inaccessible through the use of higher concentrations of the inhibitor, which proved lethal to larvae, it could be studied in *Notch1a^b420-/−^* mutants. Here all cells evidently adopted a high-Emx2 state, which drove the expression of the polarity effector above threshold and rendered them caudad-polarized (Figure 4F). Conversely, when NICD was expressed constitutively, all cells adopted a high-NICD/low-Emx2 state, inhibiting the polarity effector and becoming rostrad-polarized (Figure 4G; Supplementary Figure 10).

When Emx2 was expressed constitutively, the bistable circuit producing high- and low-NICD cells remained intact, and therefore cells assumed a low-NICD/high-Emx2 state or a high-NICD/high-Emx2 state (Figure 4H). In this case the constitutive expression of Emx2 was able to drive the expression of the polarity effector even in high-NICD cells, rendering them caudad-polarized. If NICD and Emx2 were expressed simultaneously then all cells adopted a high-NICD/high-Emx2 state in which repression of the polarity effector by NICD overcame the activation by Emx2 and produced a rostrad bias (Figure 4I). Although either treatment with DAPT or simultaneous constitutive expression of NICD and Emx2 produces a similar rostrad bias, our model predicts different mechanisms for these effects (Figure 4B; Supplementary Figure 9B). In the case of DAPT treatment we expect that most hair-cell pairs have opposite polarities, with a minority of pairs of rostrad polarity (Supplementary Information). For simultaneous constitutive expression of NICD and Emx2, however, most pairs demonstrate the same polarity, with a bias towards rostrad polarization (Supplementary information). By examining the polarity of hair cells known to be sisters, we confirmed this prediction (Supplementary Figure 11).

Our model thus explained the polarity biases observed in different experimental perturbations in terms of the cells’ joint Emx2 and NICD concentrations and the effect of these proteins on a downsteam effector of polarity reversal.

## Discussion

The coordination of polarity reversals through lateral inhibition produces pairs of oppositely polarized hair cells with virtually no errors. Our results show that robustness is ensured by the gene-regulatory network of the Notch signaling pathway, in which the downregulation of Delta ligands by NICD creates a negative feedback loop between each pair of nascent hair cells and leads to the emergence of a bistable switch (Supplementary Information) (*13*, *14*). This switch reliably breaks the symmetry between the two nascent hair cells and drives them to distinct states of Notch expression. When NICD is constitutively expressed both cells downregulate Emx2 and develop rostrad-polarized hair bundles. In *Notch1a^b420-/−^* mutants, Emx2 is upregulated in hair-cell pairs and renders them caudad-polarized. In conjunction with mathematical modelling, experiments in which NICD and Emx2 are expressed simultaneously suggest that Notch and Emx2 compete for the regulation of a polarity effector that institutes reversal of the hair-bundle orientation. Studying this effector will be the aim of future research.

To the best of our knowledge, polarity determination in the neuromast represents the first phenomenon in which the PCP and Notch pathways act in parallel to determine the orientation of a subcellular structure. The PCP pathway specifies an axis for the polarization of hair bundles; the Notch pathway then coordinates the reversal of half the bundles with respect to that axis. This process creates neuromasts in which all the cells are aligned with the anteroposterior axis of the fish, but half point in the opposite direction. This arrangement confers sensitivity to both rostrad and caudad water flows. By elucidating the interplay between cell polarity and Notch signaling, our findings explain how a complex polarity pattern is formed robustly during development and regeneration.

## Supporting information

## Acknowledgments

We thank Anna Kaczynska for expert fish husbandry and the members of our research group for comments on the manuscript. Lilianna Solnica-Krezel (Washington Univ, St. Louis) kindly provided rabbit anti-Vangl2 antiserum; Ivana Mirković and Sergiy Pylawka developed the *Tg(myo6b:GAL4FF)* line; and Sharon Amacher (Ohio State University), David Trevor (University of California, San Diego), and Kenneth Wallace (Clarkson University) provided various strains of zebrafish. A.J. was supported by an F.M. Kirby Postdoctoral Fellowship from Rockefeller University, A.E. by a Feodor Lynen Fellowship from the Alexander von Humboldt Foundation, and K.S. by Ruth L. Kirschstein National Research Service Award DC014212 from the National Institute on Deafness and Other Communication Disorders. A.D is a Postdoctoral Associate and A.J.H. an Investigator of Howard Hughes Medical Institute.

## Authors’ contributions

A.J. initiated the project, conducted experiments, developed the theoretical model, quantified the results, and wrote the paper; A.D. conducted experiments, quantified the results, and wrote the paper; A.E. conducted image analysis, quantified the results, and wrote the paper; K.S. initiated the project and conducted experiments; and A.J.H. wrote the paper.

