## Supplementary material for "Notch-Mediated Polarity Decisions in Mechanosensory Hair Cells"

**This PDF file includes:**

Materials and Methods

Theoretical Model

Figures S1 to S7

Movies S1 to S6

Tables S1 and S2

References

### Materials and Methods

*Zebrafish husbandry and strains*

Experiments were performed in accordance with the standards of Rockefeller University's Institutional Animal Care and Use Committee. Zebrafish were raised in E3 medium (5 mM NaCl, 0.17 mM KCl, 0.33 mM $CaCl_2$, 0.33 mM $MgSO_4$, 1 µg/mL methylene blue) in an incubator maintained at 28 ˚C. Wild-type TL zebrafishes were obtained from the Zebrafish International Resource Center.

The following mutant lines were used: *Notch1a*$^{-/-}$ alleles *b420* and *b638* (*1*), *Notch2*$^{-/-}$ allele *el515* (*2*), *deltaA*$^{-/-}$ allele *hi781Tg* (*3*), *deltaC*$^{-/-}$ allele *tw212b* (*4*), *deltaD*$^{-/-}$ allele *tr233Tg* (*4*), and *Emx2*$^{-/-}$ (*5*). In addition, the following transgenic zebrafish lines were used: *Tg(myo6b:actb1-EGFP)* (*6*), *Tg(myo6b:emx2-2A-NLS-mCherry)* (*5*), *Tg(5xUAS-E1b:6xMYC-notch1a-intra)* (*7*), *Tg(UAS:Tnfr2)*$^{ums1}$ (*8*), *Tg(myo6b:GAL4FF)* allele *ru1012Tg*. Embryos and larvae were staged as described (*9*).

The *Tg(myo6b:GAL4FF)* line was produced with the Gateway-based Tol2 kit (*10*). Gateway cloning was performed by combining p5E-myo6b (*6*), pME-Gal4FF (*11*), p3E-SV40-polyA (Tol2 Kit), and pDestTol2pA2 (Tol2 Kit) plasmids and LR Clonase II Plus (Invitrogen). To generate *Tg(myo6b:GAL4FF)*$^{ru1012Tg}$ founders, 25 ng/µl of verified plasmid DNA was injected into single-cell TL embryos separately from 25 ng/µl of Tol2 Transposase mRNA. Adult founders were then crossed with wild-type fish to generate heterozygotes that were identified by tail-clip genotyping with the primers: Gal4FF forward, 5′-CCAAAGAAAAACCGAAGTGC-3′; Gal4FF reverse, 5′-CTCTTCCGATGATGATGTCG-3′.

*Treatments of larvae*

To ablate mature hair cells in some experiments, we treated 3 dpf larvae for 2 hr at 28 °C with 1 µM $CuSO_4$ in E3 medium. The animals were thoroughly washed and maintained overnight in E3 medium, then fixed for immunofluorescence imaging on the following day.

After hair-cell ablation at 3 dpf, some larvae were incubated overnight with 100 μM DAPT (Cell Signaling) in E3 medium. One day later the larvae were thoroughly washed and fixed for immunofluorescence imaging.

*Immunofluorescence microscopy*

For the immunofluorescence labeling of wholemounted Emx2, Emx2-CE, and NICD-CE larvae, 4 dpf larvae were fixed overnight at 4°C in 4 % formaldehyde in phosphate-buffered saline solution (PBS) with 1 % Tween-20 (1 % PBST). Larvae were washed four times with 1 % PBST for 15 min each, then placed for 2 hrs in blocking solution containing PBS, 2 % normal donkey serum (NDS), 0.5 % Tween-20, and 1 % BSA. Primary antibodies were diluted in fresh blocking solution and incubated with the specimens overnight at 4 °C. The primary antibodies were murine anti-c-myc (1:200; Clone 9E10, Clontech), rabbit anti-Emx2 (1:200; KO609, Trans Genic, Fukuoka, Japan) (*5*), and rabbit anti-mCherry (1:200, GeneTex). Larvae were washed four times with 0.1 % PBST for 15 min each. Alexa Fluor 555/647-conjugated anti-rabbit/mouse secondary antibodies (Invitrogen, Molecular Probes), phalloidin 488 (Thermo Fisher Scientific), and 4,6′-diamidino-2-phenylindole (DAPI) were applied overnight at 1:200 dilutions in 0.2 % PBST. Larvae were then washed four times in 0.2 % PBST for 15 min each and stored at 4 °C in VectaShield (Vector Laboratories).

Immunofluorescence labeling for Delta C and Delta D was conducted on regenerating neuromasts of wholemounted larvae in which hair cells had been ablated at 3 dpf. One day later, the animals were fixed overnight at 4 °C in 4 % formaldehyde in PBS, then washed for 5 min 0.1 % PBST and placed for 2 hr in blocking solution containing PBS, 10 % NDS, 0.5 % Triton X-100, 2 % bovine serum albumin (BSA), and 1% dimethylsulfoxide (DMSO). Primary antibodies were diluted in fresh blocking solution and incubated with the specimens overnight at 4 °C. The primary antibodies were murine anti-Delta C (1:50; zdc2, Abcam) (*12*) and murine anti-Delta D (1:100; zdd2, Abcam) (*13*). Larvae were washed for 2 hr with PBS containing 0.1 % Triton X-100. AlexaFluor 647-conjugated anti-mouse secondary antibody (Invitrogen, Molecular Probes) was applied overnight at 4 °C at a dilution of 1:200 in blocking solution. Larvae were then washed 2 hr in PBS containing 0.1 % Triton X-100 and stored 4 °C in VectaShield.

Immunofluorescence labeling was used to delineate spectrin in a hair cell's cuticular plate, the actin-rich organelle at the cell's apical surface that acts as the foundation for the hair bundle. A similar procedure was employed for immunofluorescence localization of Vangl2. For wholemount preparations, 4 dpf larvae were fixed overnight at 4 °C in Prefer solution with 0.5 % Triton X-100, washed four times for 15 min each with 1 % PBST, and placed for 2 hr in blocking solution containing PBS, 2 % NDS, 0.5 % Tween-20, and 1 % BSA. Primary antibodies were diluted in fresh blocking solution and incubated with larvae overnight at 4 °C. The primary antibodies were murine anti-β-spectrin II (1:100; Clone 42/B-Spectrin II, BD Biosciences) and rabbit anti-Vangl2 (*14*). After larvae had been washed for times for 15 min each with 0.1 % PBST, we applied AlexaFluor 488- and 555-conjugated anti-mouse and anti-rabbit secondary antibodies (Invitrogen, Molecular Probes) overnight at a dilution of 1:200 in 0.2 % PBST. The specimens were then washed for times for 15 min each in 0.2 % PBST and stored 4 °C in VectaShield.

Fixed larvae were mounted on a glass slide and imaged with a microlens-based, super-resolution confocal microscope (VT-iSIM, VisiTech international) under a 100X, silicone-oil objective lens of numerical aperture 1.35. Images at successive focal depths were captured at 200 nm intervals and deconvolved with Metamorph software in ImageJ. *S*ubsequent image processing and analysis were performed with custom

ImageJ extensions written in Python. The details of the procedures and the corresponding code are provided at https://github.com/.

*Quantification of immunofluorescence images*

We developed a fully automated method for measuring the radial profile of fluorescence intensity for immunolabeled proteins at the apical surfaces of an individual hair cell. The method relies on detecting the centroid of the cell's apical surface by labeling of aconstituent of the cuticular plate such actin or spectrin. The normalized fluorescence intensity of a protein of interest, such as Vangl2 or Gnai, is then measured within a specified radial interval around the centroid for varying angles. The mean radial intensity profiles were obtained by averaging the results for 35-502 cells from 8-47 neuromasts per experimental condition,

To quantify the average fluorescence intensity of hair-cell nuclei immunolabeled for Emx2 or for NICD marked by c-myc, we combined an automated method for image segmentation with manual labelling of cells according to their polarity (Supplementary Figure 8). The region of interest within which fluorescence intensity is to be measured through a stack of images is obtained by outlining the hair cells labeled for the relevant nuclear marker. The images of individual cellular nuclei are then segmented through the stack. As an example, for a neuromast expressing β-actin-GFP under the hair-cell-specific *myo6b* promotor, the area in each optical section occupied by a particular hair cell is segmented on the basis of anti-GFP labeling. DAPI staining next allows segmentation of the nucleus within this region. The hair-bundle polarity of that cell is identified manually from the pattern of labeling with phalloidin or β-actin-GFP. Finally, a manual correction is made to the nuclear profiles to exclude any spurious area. For each nucleus, this procedure yields an average of 8-20 regions of interest from which a data point is obtained by calculating the average intensity of immunofluorescence through the entire stack and normalizing to the average intensity of DAPI staining.

To quantify the relative polarity of hair cell pairs in DAPT and NICD-CE/Emx2-CE neuromasts we manually analyzed the images and identified sister hair cells based on their similar age, size of the apical surface, and proximity. Although the identification of sister cells from fixed images is not completely unambiguous, we could nevertheless distinguish many pairs with high confidence.

### Theoretical Model

#### *Description of the model*

We consider the interaction of two cells through Notch-Delta signaling. We do not distinguish between the different Notch and Delta variants, and we disregard the role of Jagged or other ligands. The equations that describe this system are (*17*, *18*)(*15*):

$$\frac{\partial N_{1,2}}{\partial t} = N_0 H^{S_N}(I_{1,2}) - k_c N_{1,2} D_{1,2} - k_t N_{1,2} D_{2,1} - \gamma N_{1,2} \quad (1)$$

$$\frac{\partial D_{1,2}}{\partial t} = D_0 H^{S_D}(I_{1,2}) - k_c D_{1,2} N_{1,2} - k_t D_{1,2} N_{2,1} - \gamma D_{1,2} \quad (2)$$

$$\frac{\partial I_{1,2}}{\partial t} = k_t N_{1,2} D_{2,1} - \gamma_I I_{1,2} \quad (3)$$

in which $N_j$, $D_j$ and $I_j$ are the number molecules of Notch, Delta, and the Notch intra cellular domain (NICD) for cells $j = 1{,}2$. $\gamma$ represents the degradation rate constants for Notch and Delta, and $N_0$ and $D_0$ are the production rates of Notch and Delta. For simplicity we assume that both Notch and Delta degrade at a similar rate; NICD degrades at a rate $\gamma_I$. The interaction of a Notch receptor protein on one cell with a Delta ligand protein on a neighboring cell, a process called trans-activation, allows the cleavage of NICD by $\gamma$-secretase. NICD then translocates to the nucleus, where it binds to the CSL transcription-factor complex and modifies the expression of downstream targets (*16*). When the Notch receptor binds to a Delta ligand of the same cell both proteins are degraded in a process called *cis*-inhibition, and no signaling occurs. The constants $k_t$ and $k_c$ represent the constants for *trans*-activation and *cis*-inhibition respectively.

We define the Hill functions

$$H^-(I) = \frac{1}{1+(I/s_I)^n} \quad (4)$$

and

$$H^+(I) = \frac{(I/s_I)^n}{1+(I/s_I)^n} = 1 - H^-(I) \quad (5)$$

in which $s_I$ represents the binding strength of NICD to either the Notch or Delta promoters. The shifted Hill function

$$H^{S_{N,D}}(I) = H^-(I) + \lambda_{N,D} H^+(I) \quad (6)$$

represents the effect of NICD expression on the production rates of Notch and Delta where $\lambda$ is the change from the basal rate synthesis due to NICD. For $H^{S_N}(I)$ $\lambda_N > 1$, NICD activates Notch; for $H^{S_D}(I)$ $\lambda_D < 1$, NICD represses Delta. $I_0$ represents the strength of activation of Notch or inhibition of Delta by NICD.

Our experiments show that NICD downregulates Emx2 (Figures 2E-H). We describe this by introducing two equations that describe the number of Emx2 molecules $E_j$ for cells $j = 1,2$. As a simplification, supported by the temporal evolution of Delta D and Emx2 at different stages of hair-cell maturation (Figure 2D), we assume that the dynamics of Emx2 are much faster than the dynamics of the components of the Notch pathway. Because transcriptional regulation by the Notch pathway is enacted through NICD, the concentration of Emx2 depends on the concentration of NICD. Under these assumptions, we write an equation for the quasi-steady-state values of $E_j$ (*3*)

$$E_{1,2} = E_0 \frac{1}{(1+I_{1,2}/s_e)^{n_e}}, \quad (7)$$

in which $E_0$ is the steady-state number of Emx2 molecules in the absence of NICD, and $s_e$ represents the strength of the Emx2 inhibition by NICD.

Our results suggest that NICD and Emx2 compete for the regulation of a gene or set of genes that are the effectors of the polarity reversals. To test this idea we propose a simple model for polarity reversal regulation in which NICD and Emx2 act on a single downstream gene that we term polarity effector (PE). The number of molecules of this polarity effector for cells $j = 1{,}2$ is described at the quasi steady state by (*17*)

$$P_{1,2} = P_0 \frac{k_n E_{1,2}}{k_e I_{1,2} + k_n E_{1,2} + k_n k_e)}. \quad (8)$$

The NICD transcription complex binds to the polarity effector's promoter with a rate constant $k_n$ and inhibits the production of its product $P_j$. Conversely, Emx2 competes for binding at the same promoter with a rate constant $k_e$. When bound Emx2 activates the production of $P_j$. $P_0$ is the maximal steady-state number of polarity-effector molecules. For values of $P_j > P_0/2$ developing hair cells become caudad-polarized whereas if $P_j < P_0/2$, they become rostrad-polarized.

*Parameter values and robustness analysis*

The average parameter values used in simulations are described in Supplementary Table S2. The total number of proteins for Notch, Delta, and NICD were obtained from ref. (*15*). A pair of nascent hair cells remains in contact with each other for about two hours after division from their common progenitor (*18*), so we adjusted the rate constants of our model so that NICD reaches a steady state in around an hour (Figure 2A, Supplementary Figure 8). These timescales are consistent with rise times measured experimentally (*19*).

To test for robustness of the parameter values in the system, we ran 1000 simulations in which $N_0$, $D_0$, $k_c$, $k_t$, $\gamma$, $\gamma_I$, $s_I$, $s_e$, $k_n$, $k_e$, and $E_0$ to varied by 20% from the average values in Supplementary Table S2. With this level of noise in the parameters, we matched the ratio of caudad to rostrad hair cells obtained in our simulations in all of the experiments (Figure 4D-I).

*Polarity selection in wild-type neuromasts*

For the average parameter values describing a wild-type neuromast (Supplementary Table S2) the system is bistable. Starting from random initial conditions one of the two hair cells in a nascent pair will evolves to a high-Notch, high-NICD, and low-Delta state while the other cell achieves a low-Notch, low-NICD, and high-Delta state (Figure 2A, Supplementary Figure 9). If we choose cell $1$ to be in the high-Notch state, the steady state values of the variables fulfill $N_1 > N_2$, $I_1 > I_2$, and $D_1 < D_2$. NICD inhibition of Emx2 production leads to $E_2 > E_1$, and therefore $P$ is activated in cell $1$ and inhibited in cell $2$ (Supplementary Figure 8). If we choose the threshold for polarity reversals to be $P_0/2$, then cell $1$ becomes rostrad-polarized whereas cell $2$ adopts a caudad polarity.

*DAPT treatment*

DAPT is a γ-secretase inhibitor that precludes the cleavage of Notch to NICD upon binding of Notch to Delta. To analyze the effect of DAPT we need to consider a version of Eqs. (1)-(3) in which we explicitly take account of the binding step between Notch and Delta and the cleavage of DAPT. The reaction is then represented by

$$N + D_E \rightleftharpoons [ND_E] \rightarrow I \tag{9}$$

in which $D_E$ is the Delta concentration of the sister cell and $[ND_E]$ is the concentration of the uncleaved Notch-Delta complex. We can now write the equations for these reactions,

$$\frac{\partial N}{\partial t} = N_0 H^{S+}(I) - k_{t^+} N D_E - k_{t^-}[ND_E] - \gamma N, \tag{10}$$

$$\frac{\partial [ND_E]}{\partial t} = -k_{t^+} N D_E - k_{t^-}[ND_E] - k_I[ND_E], \tag{11}$$

$$\frac{\partial I}{\partial t} = -k_I[ND_E] - \gamma_I I, \tag{12}$$

in which $k_{t^+}$ and $k_{t^-}$ represent the binding and unbinding rate constants of Notch to Delta and $k_I$ is the cleavage rate constant for the Notch-Delta complex. For simplicity we have omitted the hair-cell subscripts, the terms describing *cis*-inhibition, and the equation for Delta.

Assuming a quasi-steady state for $[ND_E]$, from Eq. (11) we obtain

$$[ND_E] = \frac{k_{t^+}}{k_{t^-} + k_I} ND_E. \tag{12}$$

Inserting Eq. (12) into Eqs. (10) and (12), we obtain

$$\frac{\partial N}{\partial t} = N_0 H^{S+}(I) - k_t N D_E - \gamma N, \tag{13}$$

$$\frac{\partial I}{\partial t} = -k_t N D_E - \gamma_I I, \tag{14}$$

with

$$k_t = \frac{k_I k_{t^+}}{k_I + k_{t^-}}. \tag{15}$$

Addition of DAPT lowers the cleavage rate constant $k_I$ of the Notch-Delta complex, which implies a decrease of $k_t$. Therefore, decreasing values of $k_t$ in Eqs. (1)-(3) simulate the effect of increasing concentrations of DAPT (Supplementary Figure 9). For a moderate decrease of $k_t$ the concentration of NICD for the high-NICD cell decreases but the system retains its bistability and the polarity effector $P$ is driven to a high state, which induces a caudad polarization of the cell. For higher concentrations of DAPT, corresponding to a lower value of $k_t$, a new, symmetric, state appears as a middle branch of the bifurcation diagram (Supplementary Figure 9C). Both cells in this state show intermediate to low concentrations of NICD. For $k_t \gtrapprox 0.2 \times 10^{-6} s^{-1}$, the NICD expression of this symmetric state is sufficient to repress the expression of Emx2 and the polarity effector to produce pairs of rostrad-polarized cells. As a result, when we introduce noise in the parameters of the system while using a low value of $k_t$ to simulate a DAPT treatment many cell pairs adopt this intermediate state, generating the rostrad bias observed experimentally (Supplementary Table S2, Figure 4E, Supplementary Figure 9).

For $k_t \lessapprox 0.2 \times 10^{-6} s^{-1}$ our model predicts that the NICD concentration of the symmetric state is insufficient to repress Emx2 and the polarity effector, and therefore the cells become caudad-polarized. This regime corresponds to high concentrations of DAPT that are unfortunately lethal to the fish.

*Notch 1a mutants*

In Notch 1a null mutants, the rate of production of functional Notch is $N_0 = 0\ s^{-1}$. This case is equivalent to the limit in which $k_t = 0$. In the complete absence of NICD, Emx2 is expressed highly in both cells of the pair, activating the expression of the polarity effector and creating caudad-polarized cells (Figure 2F).

*Constitutive expression of NICD*

The constitutive expression of NICD under the myo6b promoter can be simulated by adding a source term $I_0$ to Eq. (3), which represents the extra rate of production of NICD

$$\frac{\partial I_{1,2}}{\partial t} = k_t N_{1,2} D_{2,1} - \gamma_I I_{1,2} + I_0. \tag{16}$$

The bifurcation diagram of the system as a function of $I_0$ shows that moderate to high rates of constitutive NICD production inhibit the expression of Emx2, as observed experimentally (Figure 2H, Supplementary Figure 10). The higher repression by NICD and lower activation by Emx2 lead to below-threshold levels of the polarity effector, producing a bias towards rostrad-polarized cells (Figure 2G, Figure 4G, Supplementary Figure 10). Above a certain value of $I_0$ the bistable steady state of the wild-type system vanishes and both cells of a nascent pair adopt intermediate to high levels of NICD expression. For the robustness analysis of this case we allow for the variation of all the parameter values described before as well as of $I_0$ (Figure 4G).

*Constitutive expression of Emx2*

The constitutive expression of Emx2 increases the concentration of this protein at the steady state. To simulate this case we add a constant term $E_c$ to Eq. (7):

$$E_{1,2} = E_0 \frac{1}{(1 + I_{1,2}/s_e)^{n_e}} + E_c. \tag{17}$$

Because there is no feedback from Emx2 to the Notch signaling pathway, the constitutive expression of Emx2 does not affect the expression of NICD. Therefore, the bistable state with high-NICD and low-NICD persists (Figure 4H, Supplementary Figure 6). In the low-NICD cell the concentration of Emx2 is even higher than in a low-NICD hair cell of a wild type larva and this cell remains caudad-polarized. For the high NICD cell, the increased concentration of Emx2 suffices to counteract the inhibition of the polarity factor by NICD, and exeeds the threshold to produce caudad-polarized cells (Figure 4H). Despite the presence of cells with two distinct NICD states, the constitutive expression of Emx2 therefore specifies polarity reversal (Supplementary Figure 6). For the robustness analysis of this case we allow for the variation of all the parameter values described before as well as for $E_C$ (Figure 4H)

*Simultaneous constitutive expression of NICD and Emx2*

We simulate the simultaneous constitutive expression of NICD and Emx2 by combining Eqs (16) and (17). The constitutive expression of NICD results in the disappearance of the bistable state seen in wild-type larvae (Supplementary Figure 8, Supplementary Figure 10) and both cells of the pair adopt a high-NICD state (Figure 2G). At the same time, because of the constitutive expression of Emx2, both cells achieve high Emx2 levels despite the high NICD concentrations. A tug-of-war then ensues between NICD and Emx2 for the activation or inhibition of the polarity factor. We experimentally observe that simultaneous constitutive expression of NICD and Emx2 leads to a rostrad bias (Figure 4A-B). This suggests that either NICD is constitutively expressed at much higher levels than Emx2, or that NICD more effectively inhibits the polarity effector than Emx2 activates it (Figure 4I).

#### *Sources of polarity bias in DAPT-treated and NICD-CE/Emx2-CE neuromasts*

Neuromasts of DAPT treated larvae, and those with simultaneous constitutive expression of NICD and Emx2 show a similar rostrad bias (Figure 4B, Supplementary Figure 9B). Our model shows that the origin of this bias has a different origin for each of these two conditions. For DAPT treated neuromasts the hair cell pairs can either adopt an asymmetric state, with a high- and low-NICD cells, or a symmetric state, with intermediate-NICD levels for both cells (Supplementary Figure 9C). Cells pairs in the asymmetric state will adopt opposite polarities while those in the symmetric state will adopt equal polarities.

In the case of NICD-CE/Emx2-CE larvae, most cells pairs will adopt a symmetric high-NICD state (Supplementary Figure 10), and therefore most sister cell pairs will show the same polarity. Fluctuations in the level of NICD and Emx2 for these pairs will determine if both cells are caudad or rostrad polarized.

To test this prediction of our model we identified sister cell pairs in DAPT and NICD-CE/Emx2-CE images and quantified the ratio of same polarity pairs (Supplementary Figure 11). Our data confirms our theoretical prediction and shows that in DAPT treated neuromasts only around 50% of hair cell pairs show opposite polarity, compared to about 70% in NICD-CE/Emx2-CE neuromasts.

### Supplementary figures

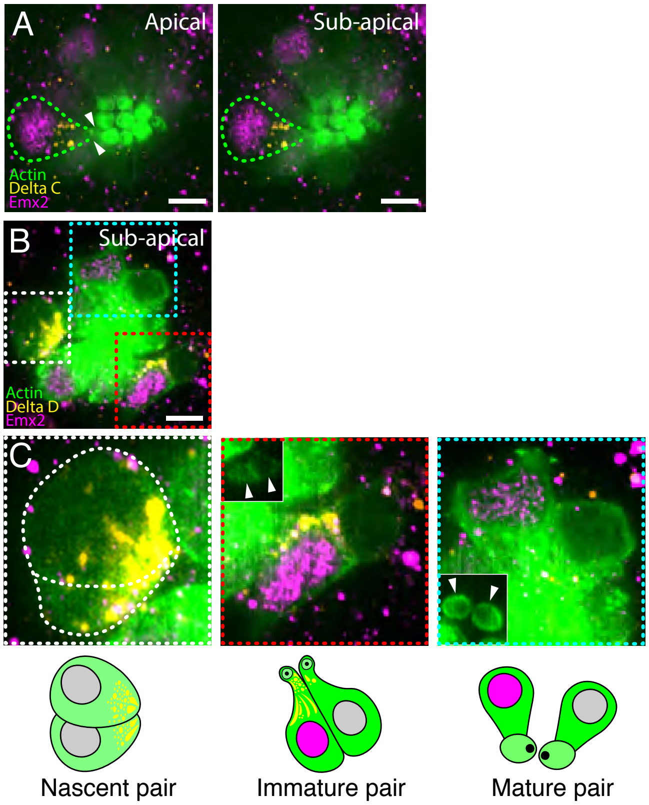


**Figure S1. Expression of Delta C and Delta D at successive stages of hair-cell development.** (**A**) In apical and sub-apical views of a 4 dpf neuromast GFP-labeled actin (green) marks hair cells. One hair cell of an immature pair (arrowheads) expresses Emx2 (magenta) and Delta C (orange); its soma is outlined. (**B**) A sub-apical view of a 4 dpf neuromast shows pair of hair cells at the nascent (white box), immature (red box), and mature stages (cyan box). GFP-labeled actin (green) marks hair cells; Emx2 (magenta) and Delta D (orange) are immunolabeled. (**C**) Enlarged views of hair cells at the three stages (above) are paired with schematic diagrams (below) of the different stages. The insets depict the apical surfaces of the corresponding hair-cell pairs (arrowheads). Scale bars, 5 $\mu$m.

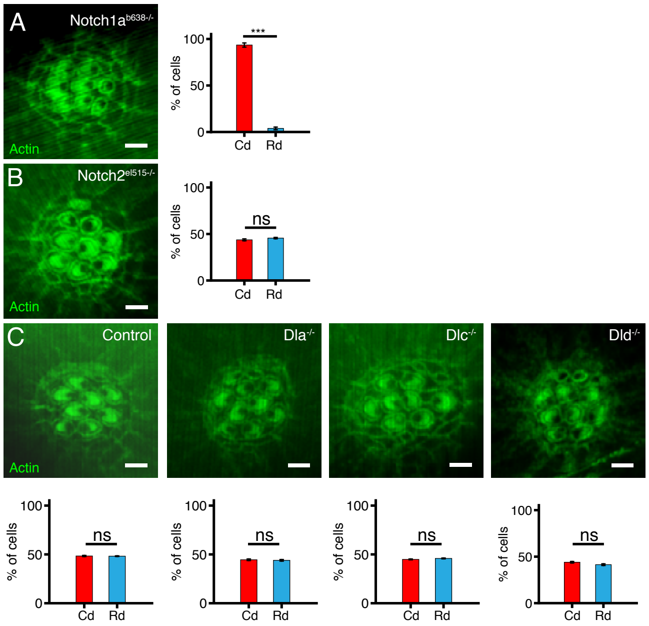


**Figure S2. Hair-cell polarity in *Notch1a$^{b638-/-}$*, *Notch2$^{el515-/-}$*, and Delta mutant neuromasts.** (**A**) Phalloidin-stained actin (green) marks the apical surfaces of hair cells in a 4 dpf *Notch1a$^{b638-/-}$* larva. The histogram quantifies the percentages of hair cells of caudad (Cd) and rostrad (Rd) polarities in 58 hair cells from 11 neuromasts. (**B**) The corresponding micrograph and histogram for a 4 dpf *Notch2$^{el515-/-}$* larva show no polarity bias in 276 hair cells from 31 neuromasts. (**C**) The corresponding data from 4 dpf neuromasts of wild-type, *Dla$^{-/-}$*, *Dlc$^{-/-}$*, and *Dld$^{-/-}$* larvae again reveal no polarity bias. The results represent 292 hair cells from 38 neuromasts for *Dla$^{-/-}$*, 200 hair cells from 19 neuromasts for *Dlc$^{-/-}$*, and 120 hair cells from 14 neuromasts for *Dld$^{-/-}$*. Scale bars, 2 $\mu$m. Means ± SDs; ns = not significant.

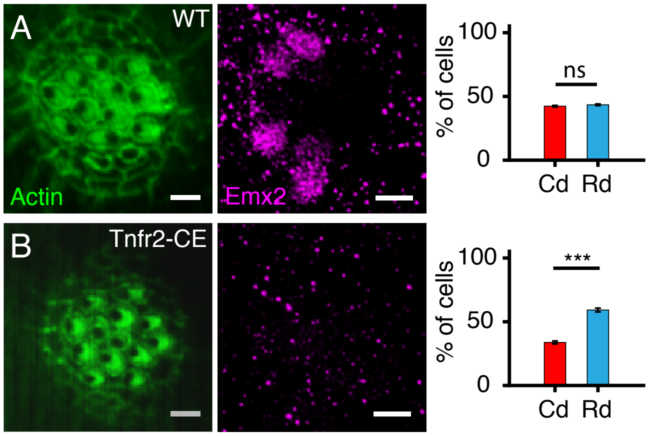


**Figure S3. Rostrad polarity bias from constitutive expression of tumor necrosis factor 2 (Tnfr2-CE).** (**A**) In an apical view of a 4 dpf wild-type neuromast, phalloidin labels actin (green) and Emx2 is immunolabeled (magenta). The associated histogram shows no polarity bias between hair cells of caudad (Cd) and rostrad (Rd) polarity. (**B**) A corsponding view of a 4 dpf Tnfr-CE neuromast shows a propondeance of rostrad-polarized hair cells. The histogram confirms this asymmetry for 471 hair cells from 40 neuromasts. Scale bars, 2 $\mu$m. Means ± SDs; ns = not significant; *** = $p < 0.001$.

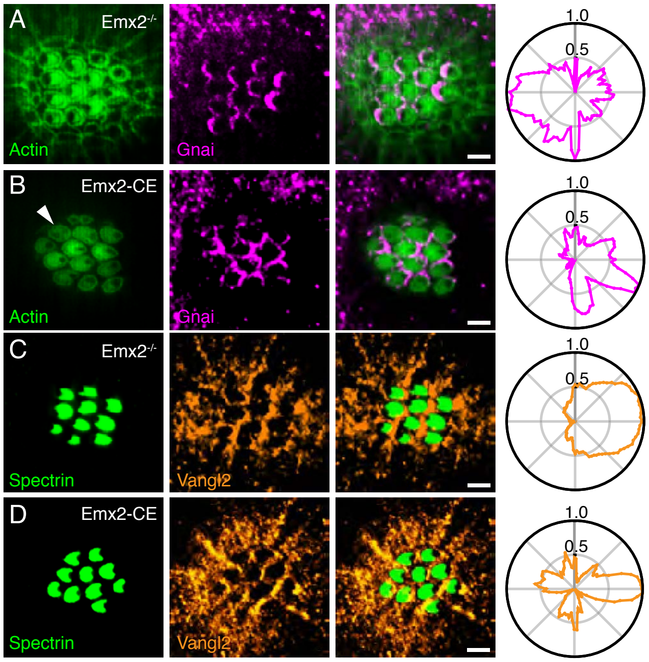


**Figure S4. Expression of Gnai and Vangl2 in *Emx2*$^{-/-}$ and Emx2-CE neuromasts.** (**A**) In an apical view of a 4 dpf *Emx2*$^{-/-}$ mutant neuromast, hair cells marked by actin-GFP (green) show a uniformly rostrad orientation. Immunolabeled Gnai (magenta) occurs at the anterior edges of the cells, adjacent to the kinocilia. The histogram confirms this asymmetry for 90 hair cells. (**B**) In an apical view of a 4 dpf *Emx2*$^{-/-}$ mutant neuromast immunolabeling for spectrin (green) reveals a uniformly rostrad orientation of the hair cells. Immunolabeling for Vangl2 (orange) indicates that this PCP protein occurs uniformly at the posterior edges of hair cells, as is confirmed by a histogram for 165 hair cells. (**C**) In an apical view of a 4 dpf Emx2-CE neuromast, hair cells marked by actin-GFP (green) show a rostrad orientation. The polarity of a single hair cell is ambiguous (arrowhead). Immunolabeled Gnai (magenta) occurs at the posterior edges of the cells, adjacent to the kinocilia. The histogram confirms this orientation for 36 hair cells. (**D**) In an apical view of a 4 dpf Emx2-CE neuromast, immunolabeling for spectrin (green) reveals a uniformly caudad orientation of the hair cells. Immunolabeled Vangl2 (orange) again occurs consistently at the posterior edges of hair cells, a pattern confirmed by a histogram from 163 hair cells. Scale bars, 2 $\mu$m.

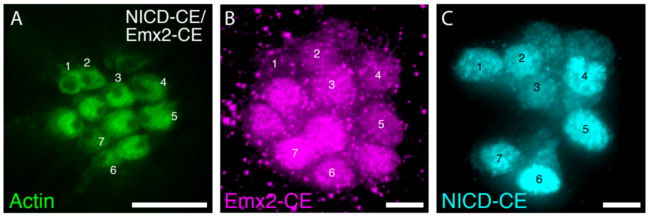


**Figure S5. Rostrad polarity bias from simultaneous constitutive expression of NICD and Emx2.** (**A**) In an apical view of a 4 dpf NICD-CE/Emx2-CE neuromast, GFP-labeled actin (green) marks hair cells. (**B**) mCherry immunolabeling (magenta) denotes the hair cells that constitutively express Emx2. (**C**) Immunostaining for NICD-myc (cyan) reveals the hair cells that expressing NICD. Numbers indicate mature hair cells that express both NICD (cyan) and Emx2 (magenta). Note that these seven cells express manifest a rostrad orientation, as opposed to the caudad polarity of the other mature hair cells. Two immature hair cells occur at the top of the neuromast. Scale bars, 5 $\mu$m.

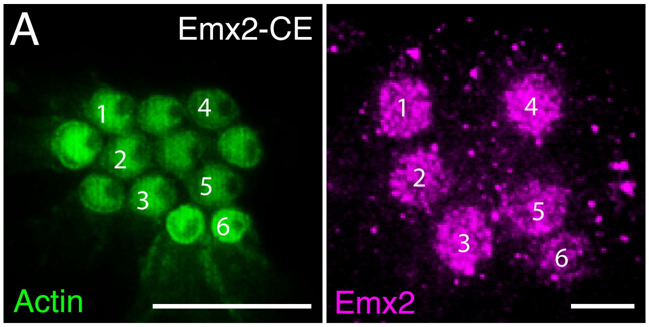


**Figure S6. Emx2 bistability is maintained in hair cells constitutively expressing Emx2.** (A) In an apical view of a 4 dpf Emx2-CE neuromast, hair cells marked by actin-GFP (green) show a uniformly caudad orientation. Emx2 bistability, as revealed by immunolabeling (magenta, numerals), is maintained irrespective of constitutive expression of Emx2.

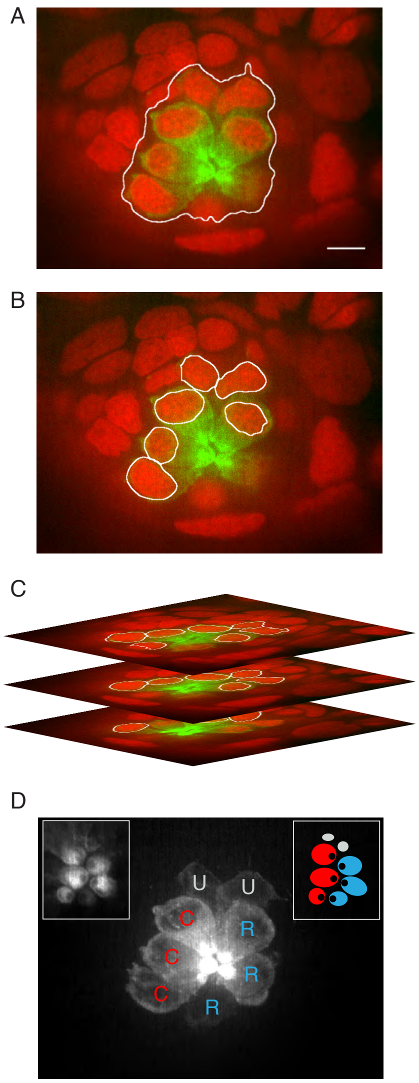


**Figure S7. Quantification of relative nuclear immunofluorescence.** (**A**) In a *Tg(myo6b:bactin-GFP)* neuromast, GFP (green) marks actin and DAPI (red) stains nuclei. Automated segmentation defines the region containing the GFP signal (outline). (**B**) The nuclei are subsequently segmented on the basis of DAPI staining. (**C**) An average of 8-20 regions of interest are obtained for each hair cell in a through-focus series. The immunofluorescence signal for the marker of interest, Emx2 or NICD, is averaged across these regions of interest. (**D**) In a manual step, polarity of each hair cell is identified from actin-GFP labeling (white), which marks the hair cells and their bundles (insets). The insets portray the apical hair-cell surfaces (left) and a diagram of the hair-bundle polarities (right). C, caudad; R, rostrad; U, unidentified. Scale bar, 5 $\mu$m.

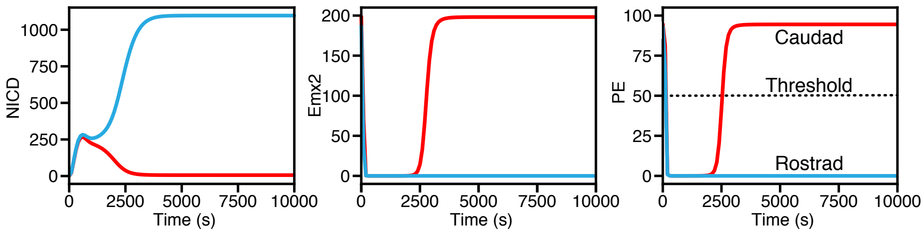


**Figure S8. Simulation of wild-type responses in the polarity-effector model.** (**A**) The panels show time traces for NICD (left), Emx2 (center), and the polarity effector (PE, right) in a simulation of Equations 1-3 (Supplementary Material). The parameter values correspond to those for a wild-type neuromast (Supplementary Table 2). The high-NICD receiver cell (blue line) ceases to produce Emx2, has a negligible concentration of PE, and therefore adopts a rostrad polarity. The low-NICD sender cell (red line) attains a high concentration of Emx2, activates the expression of PE to a level exceeding the threshold, and thus acquires a caudad polarity.

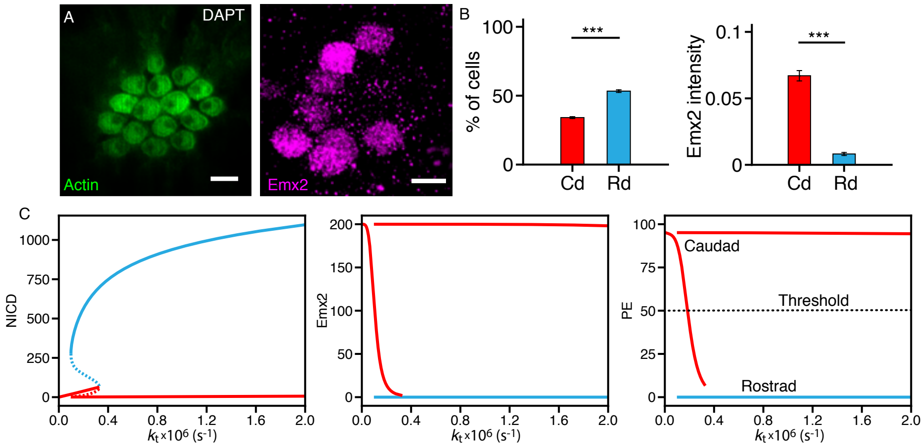


**Figure S9. Rostrad polarity bias from Notch inhibition by DAPT.** (**A**) In an apical view of a 4 dpf DAPT-treated neuromast. GFP-labeled actin (green) marks hair cells and Emx2 is immunolabeled (magenta). (**B**) The polarities of 175 hair cells from 31 DAPT treated neuromasts (left) show a bias toward rostrad polarization. The average level of Emx2 expression (right) is similar to wild-type larvae. (**C**) Bifurcation diagrams show the number of molecules of NICD (left), Emx2 (center), and polarity effector (PE, right) as a function of the Notch cleavage rate ($k_t$). Decreasing values of this rate correspond to increasing values of the inhibitor DAPT. Without DAPT ($k_t = 2 \times 10^{-6}$ s$^{-1}$), one cell adopts a high-NICD state and rostrad polarity (blue line) and the other a low NICD state and caudad polarity (red line). For increasing concentrations of DAPT the concentration of NICD in the high expressing cell decreases but not enough to change its polarity. For high concentrations of DAPT ($k_t \approx 3 \times 10^{-6}$ s$^{-1}$), a new state emerges in which both cells express intermediate NICD concentrations. Although this state yields rostrad-polarized pairs of hair cells for moderate NICD concentrations, for even higher concentrations it would produce caudad-polarized pairs. Scale bars, 5 $\mu$m. *** = $p < 0.001$.

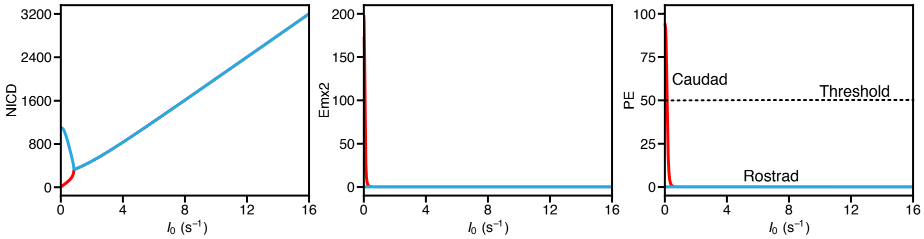


**Figure S10. Caudad polarity bias from constitutive NICD expression.** (**A**) A bifurcation diagram shows the number of molecules of NICD (left), Emx2 (center), and polarity effector (PE, right) as a function of the rate of constitutive expression of NICD ($I_0$). Without NICD ($I_0 = 0$), one hair cell adopts a high NICD-state and rostrad polarity (blue line) and the other a low-NICD state and caudad polarity (red line). As the rate of of constitutive expression of NICD increases, however, the bistable state is replaced by a symmetric state of high NICD concentration. In this state both cells adopt a rostrad polarity.

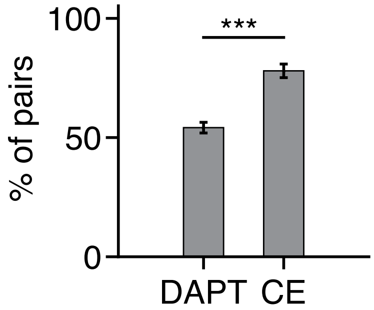


**Figure S11. Distinct sources of rostrad bias for simultaneous expression of NICD and Emx2 *versus* DAPT treatment.** The histogram shows the percentages of sister hair cells of the same polarity following DAPT treatment in 153 cell pairs of 27 neuromasts and in 38 cell pairs of 33 NICD-CE/Emx2-CE neuromasts. In DAPT treated neuromasts a many hair cell pairs adopt a bistable steady state and therefore opposite polarizations (Supplementary Figure 9). For simultaneous constitutive expression of NICD and Emx2 most cells adopt a symmetric, high Notch, state, and therefore have the same polarization (Supplementary Figure 10).

**Supplementary movies**

**Movie S1.** In a through-focus scan of a 4 dpf *Tg(myo6b:bactin-GFP)* neuromast, hair cells are labeled green with β-actin-GFP. Three mature hair cells express Emx2 (magenta); the immature hair cell at the top expresses both Emx2 and Delta D (orange). Scale bar, 5 µm.

**Movie S2.** In a through-focus scan of a 4 dpf *Tg(myo6b:bactin-GFP)* neuromast, hair cells are labeled green with β-actin-GFP. Four mature hair cells express Emx2 (magenta), whereas the sister cells of opposite polarity do not. Immature hair cells on the left and right sides of the neuromast express both Emx2 and Delta C (orange). Scale bar, 5 µm.

**Movie S3.** In a through-focus scan of a 4 dpf *Tg(myo6b:bactin-GFP)* neuromast, hair cells are labeled green with β-actin-GFP. Four mature hair cells express Emx2 (magenta), whereas two do not. One hair cell of the immature at the lower right expresses both Emx2 and Delta D (orange); its sister expresses neither. Note the Delta D-labeled nascent hair cells on the left. Scale bar, 5 µm.

**Movie S4.** In a through-focus scan of a 4 dpf *Tg(myo6b:bactin-GFP); Tg(myo6b:GAL4FF); Tg(5xUAS-E1b:6xMYC-notch1a-intra)* neuromast, hair cells are labeled green with β-actin-GFP. Seven of the mature hair cells constitutively express NICD (cyan) and display rostrad polarity. Two mature hair cells contain Emx2 (magenta), however, and show caudad sensitivity. Scale bar, 5 µm.

**Movie S5.** In a through-focus scan of a 4 dpf *Notch1a*$^{b420-/-}$ neuromast, the apical surfaces of hair cells are stained green with phalloidin. All ten hair cells express Emx2 (magenta) and show a caudad polarization. Scale bar, 5 µm.

**Movie S6.** In a through-focus scan of a 4 dpf NICD-CE/Emx2-CE neuromast, hair cells are labeled green with β-actin-GFP. The apical surfaces (left) reveal that four of the mature hair cells show caudad polarization and seven display rostrad sensitivity. Although all eleven of these cells express mCherry (magenta, center), those of a rostrad polarity also express NICD (cyan, right). Scale bar, 5 μm.

### Supplementary tables

#### Table S1: Abbreviations of transgenic constructs

| Abbreviations | Genetic background | Details |
|---|---|---|
| NICD-CE | *Tg(myo6b:actb1-EGFP); Tg(myo6b:GAL4FF); Tg(5xUAS-E1b:6xMYC-notch1a-intra)* triply transgenic zebrafish | β-actin-EGFP marks hair cells; Notch1a intra-cellular domain tagged with c myc is constitutively expressed in hair cells. |
| Emx2-CE | *Tg(myo6b:actb1-EGFP); Tg(myo6b:emx2-2A-NLS-mCherry)* doubly transgenic zebrafish | β-actin-EGFP marks hair cells; full-length Emx2 is constitutively expressed in hair cells; mCherry expression in the nucleus marks hair cells constitutively expressing Emx2. |
| NICD-CE/Emx2-CE | *Tg(myo6b:actb1-EGFP); Tg(myo6b:GAL4FF); Tg(5xUAS-E1b:6xMYC-notch1a-intra); Tg(myo6b:emx2-2A-NLS-mCherry)* quadruply transgenic zebrafish. | β-actin-EGFP marks hair cells; both NICD and Emx2 are constitutively expressed in hair cells; anti-c-myc (cyan) and anti-mCherry (magenta) antibodies mark NICD+Emx doubly positive hair cells. |
| Tnfr2-CE | *Tg(myo6b:GAL4FF); Tg(UAS:Tnfr2)* doubly transgenic zebrafish. | Tumor necrosis factor 2 (Tnfr2) is constitutively and specifically expressed in hair cells. |

**Table S2: Values of model parameters**

| Parameter | Mean value | Unit |
|---|---|---|
| $\gamma$ | $1\times10^{-3}$ | time$^{-1}$ (s$^{-1}$) |
| $\gamma_I$ | $5\times10^{-3}$ | time$^{-1}$ (s$^{-1}$) |
| $k_t$ | $2\times10^{-6}$, $3\times10^{-7}$ * | time$^{-1}$ (s$^{-1}$) |
| $k_c$ | $5\times10^{-6}$ | time$^{-1}$ (s$^{-1}$) |
| $N_0$ | 5, 0 ** | time$^{-1}$ (s$^{-1}$) |
| $D_0$ | 10 | time$^{-1}$ (s$^{-1}$) |
| $s_I$ | 200 | Dimensionless |
| $n$ | 2 | Dimensionless |
| $\lambda_N$ | 2 | Dimensionless |
| $\lambda_D$ | 0 | Dimensionless |
| $E_0$ | 200 | Number of proteins |
| $s_e$ | 20 | Number of proteins |
| $n_e$ | 4 | Dimensionless |
| $P_0$ | 100 | Number of proteins |
| $k_n$ | 40 | Dimensionless |
| $k_e$ | 10 | Dimensionless |
| $I_0$ | 16 *** | time$^{-1}$ (s$^{-1}$) |
| $E_c$ | 200 **** | Number of proteins |

* DAPT treatment (Figure 2E)
** Notch 1a mutant (Figure 2F)
*** Constitutive expression of NICD (Figure 2G,I)
**** Constitutive expression of Emx2 (Figure 2H,I)
